# Prophage rates in the human microbiome vary by body site and host health

**DOI:** 10.1101/2023.05.04.539508

**Authors:** Laura K. Inglis, Michael J. Roach, Robert A. Edwards

## Abstract

Phages integrated into a bacterial genome–called prophages–continuously monitor the health of the host bacteria to determine when to escape the genome, protect their host from other phage infections, and may provide genes that promote bacterial growth. Prophages are essential to almost all microbiomes, including the human microbiome. However, most human microbiome studies focus on bacteria, ignoring free and integrated phages, so we know little about how these prophages affect the human microbiome. We compared the prophages identified in 11,513 bacterial genomes isolated from human body sites to characterise prophage DNA in the human microbiome. Here, we show that prophage DNA comprised an average of 1-5% of each bacterial genome. The prophage content per genome varies with the isolation site on the human body, the health of the human, and whether the disease was symptomatic. The presence of prophages promotes bacterial growth and sculpts the microbiome. However, the disparities caused by prophages vary throughout the body.

## Introduction

The human microbiome is a complex ecosystem of microbes that inhabit every part of the human body. Most body sites typically contain a multitude of microbes resulting in diverse ecosystems. In contrast, sites in the human body dominated by one or a few species—dysbiosis—are often an indicator of disease ^1–3^; Inglis and Edwards 2022).

While the term ‘human microbiome’ may evoke the mental image of the human body as a single environment, the body contains many different niches. Environments such as the skin, stomach, lungs, and mouth are so different from each other that combined, they have an extensive range of bacterial concentrations, very high species richness, and widely varying species diversities. Most of the total microbial biomass in humans and other mammals resides in the gut, and that organ’s metabolism contributes to the animal’s overall thermogenic energy expenditure ^4^. Other body areas have orders of magnitude lower bacterial concentrations than the gut ^5^. The gut microbiome is also highly diverse ^6^, while others, such as the lung microbiome, are dominated by only a few groups ^7^.

Bacteriophages (phages) are viruses that infect bacteria found in almost every environment ^8^. In the ocean, they kill around 20% of the microbial biomass daily ^9^, but their role in sculpting and controlling most microbiomes, including the human microbiome, is underestimated. There are two main kinds of phages: virulent phages, where the phage infects the host bacteria, replicates, and lysis the bacteria to release phage progeny, and temperate phages, which may either choose a lytic lifecycle or choose to integrate into the host’s DNA to be passively replicated alongside the host until the phage senses suitable conditions for the switch to lytic replication ^10^. Prophages are temperate phages integrated into their host’s genome, and the resulting host bacteria is a ‘lysogen’. Almost every bacterial species has temperate phages, although much is still unknown about both lytic and temperate phages.

Prophages confer various benefits to their host through lysogenic conversion. The most common is superinfection exclusion: the protection of the lysogen against other phage infections ^11,12^. Many prophages also express virulence genes or toxins that promote the growth of the lysogen ^13–17^2; von Wintersdorff et al. 2016; Waldor and Mekalanos 1996). Some examples of prophages providing the toxins that allow their bacterial host to cause human disease include Shiga toxin-producing *E. coli*, cholera and diphtheria.

The genetic switch that controls the decision to integrate into the host or replicate and kill the host has been at the centre of many molecular biology breakthroughs ^18^, such as the Nobel prizes in physiology and medicine in 1965 and 1969 which were awarded for discoveries regarding the viral synthesis and replication mechanisms respectively.

Many factors affect the outcome of that decision, including the concentration of bacteria, the diversity of bacterial and phage species, the redox potential of the cell (i.e. the metabolic efficiency of the bacteria), the presence of other phages, and signalling peptides that phages produce to communicate with each other ^19,20^.

Here we explored the variation in prophage composition across the human body, investigated how much of the bacterial DNA in the human microbiome is provided by prophages, demonstrated how this diverges across the different areas of the human body, and we quantified whether diseased microbiomes and disease-causing bacteria have different prophage abundances than the microbiomes of healthy people.

## Results and discussion

The GenBank genome assembly database contains almost 1 million publicly accessible bacterial genomes, but most are highly fragmented. However, we identified 11,513 genomes from bacteria that could be associated with different areas of the human body. These samples came from various people with different geographical locations, lifestyles, ages, diets, and conditions. We identified prophages in these genomes and calculated the percentage of the prophage sequence genomes for each sample source location (Fig. 1).

**Figure 1:**
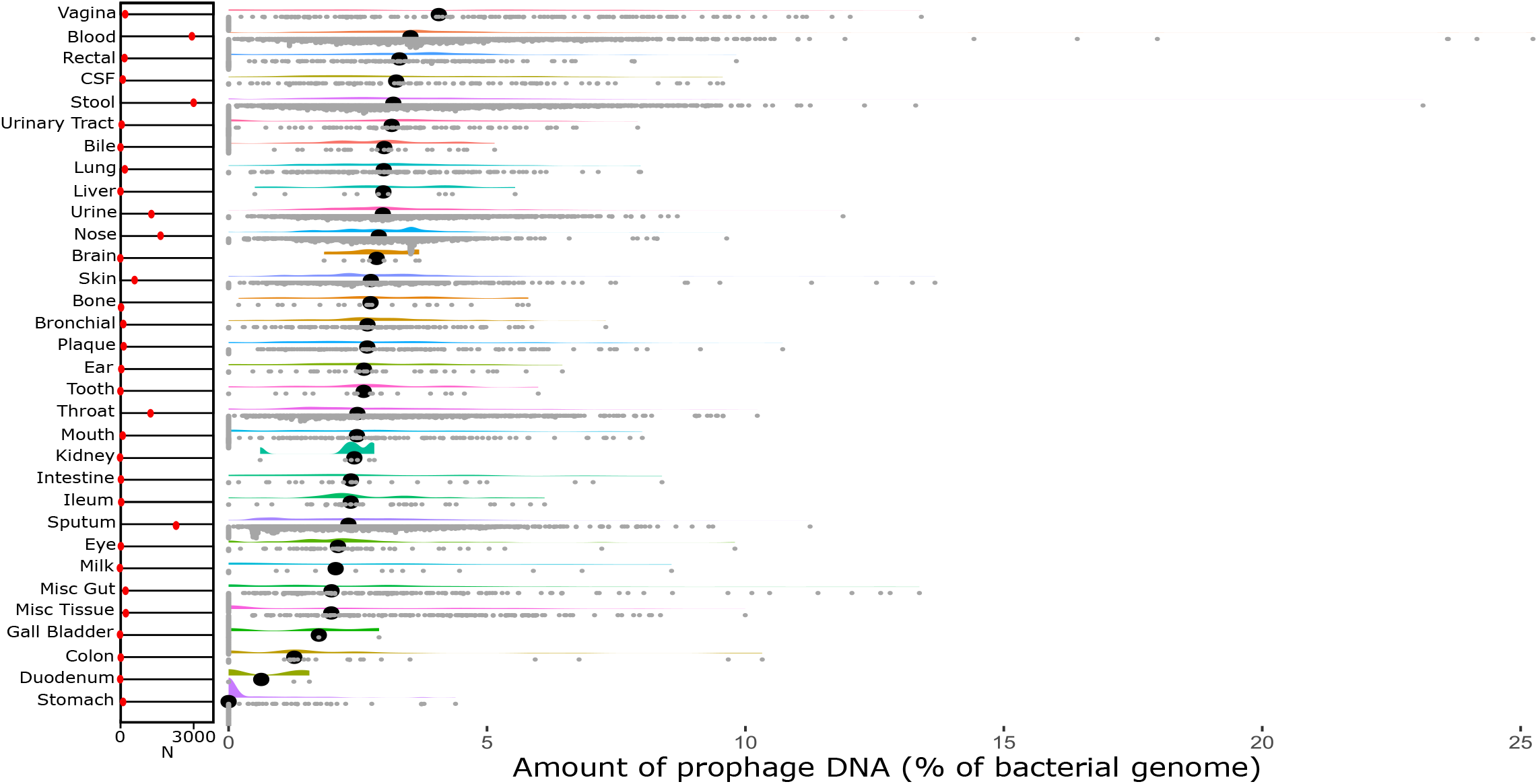
Raincloud plot of all 32 categories ordered by the median percentage of phage DNA from highest to lowest. The red markers on the left show the number of genomes in each category.

While the areas had different amounts of genomes associated with them (Fig 1; red markers), almost every area had a considerable variation in the number of prophages. This variation could primarily be due to the differences in microbiome bacterial compositions between the individuals sampled. While there is evidence for a ‘core microbiome’ of functional genes ^21–23^, the taxonomic makeup of the microbes between individuals varies significantly ^24^. Many factors affect the composition of our microbiomes, including diet, medications, overall health and fitness, and weight ^1,2,21,25–27^.

There is a large variation in the proportion of prophage DNA within and between different body sites. The median prophage DNA content ranges from 0-5% of the bacterial genome. While many areas have a median prophage DNA content closer to 2-3%, there is a sizeable difference between body sites, especially at the extremes--vagina and blood at the high end, with 4-5% prophage content, and duodenum, and stomach at the lower end, with close to 0% prophage content.

The vaginal samples had the highest median proportion of prophage DNA. A single genus of bacteria dominates the healthy vaginal microbiome—*Lactobacillus*—which produces antimicrobial compounds that control other bacterial populations ^28^. The vaginal microbiome is also dense, containing 10^10^-10^11^ bacterial cells ^28^. Both high bacterial concentrations and a microbiome dominated by a few species are two factors previously shown to correlate with higher rates of lysogeny ^20^.

Conversely, the stomach had the lowest average proportion of prophage DNA. No prophages could be detected in most (76.67%) of the genomes from bacteria isolated in the stomach. The stomach is significantly different from almost every other body site and is one of the most extreme environments in the human body. A handful of genera dominate, and the bacterial concentrations are relatively low, in the order of 10^3^-10^4^ bacteria ^29^. Overall, It is quite the opposite of the vaginal microbiome.

## Respiratory and gastrointestinal tracts

Narrowing our focus to the respiratory and gastrointestinal tracts allows us to examine how the microbiome changes as the conditions change in transit from mouth to anus. There are bacteria everywhere; some enter our bodies through our mouth/nose as we eat, drink and breathe, and some of these bacteria find their way down further into the respiratory or gastrointestinal systems to supplement those microbiomes ^30^. This results in connected microbiomes, such as the mouth, nose, and lungs, having similar microbiome compositions. We juxtaposed the distributions of prophage DNA with the GI and respiratory systems for visual assessment (Fig 2), and we performed Kruskal-Wallis statistical tests to determine if these distributions were significantly different.

**Figure 2:**
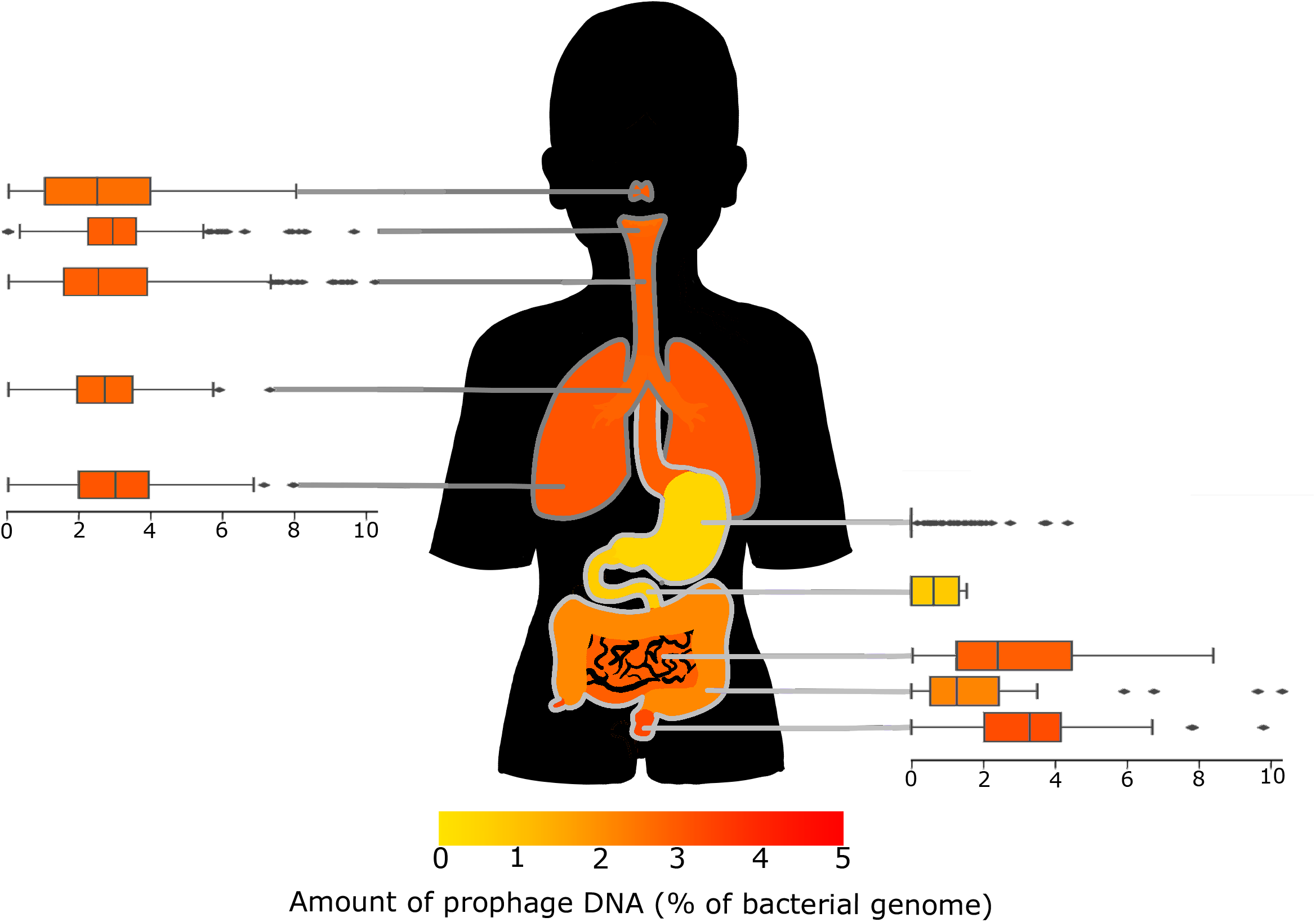
Box plots showing the amount of prophage DNA in each area of the respiratory and gastrointestinal tracts. The figure is coloured by the average proportion of prophage DNA as displayed in the scale below.

The different sections of the respiratory tract have similar distributions of prophage DNA, reflecting similar conditions. The areas connected to the gastrointestinal tract--the mouth and throat--were significantly different to the lungs (p=<0.05), while the nose was only significantly different to the throat (p=<0.005).

The lungs had the highest prophage DNA, while the mouth had the lowest. The microbiome of the respiratory tract changes with the age of the host, becoming more diverse as the human matures from infant to adult ^31^. Since the abundance of temperate phages correlates with microbial diversity, there may be fewer temperate phages in the respiratory tract of older people. The overall bacterial concentration estimates suggest the mouth has more bacteria than the lungs ^5,30^, which is generally conducive to higher rates of lysogeny. However, we observe the opposite trend with lung-isolated genomes having higher proportions of prophage DNA.

There were two main hypotheses regarding lysogeny rates, piggyback-the-winner and piggyback-the-persistent. Piggyback-the-winner suggests that the microbiomes with high bacterial concentrations are more likely to favour lysogeny ^32^, while the piggyback-the-persistent suggests the opposite ^33^. The lungs have a lower bacterial concentration, yet a relatively high amount of prophage DNA suggests that it might follow the persistent strategy of piggyback.

Conversely, the gastrointestinal tract has a much wider range of bacterial concentrations and does not follow a linear order like the respiratory tract. The distinct areas of the gastrointestinal tract have much more varied environments, and the prophages appear to follow the Piggyback-the-winner model, with more prophages in areas of the body with higher bacterial concentrations, such as the stool or mouth. In contrast, the more hostile environments like the stomach have less prophage DNA per bacteria.

Overall, bacterial concentration alone does not adequately explain the proportion of prophage DNA, and we must look to other factors to explain our results. Generally, bodily fluid samples (e.g. breast milk, urine, and blood) had lower bacterial concentrations ^34–37^ but higher prophage concentrations than the other body sites.

### Effects of host health

Because different bacteria have different prophages and lower bacterial diversity is associated with higher rates of lysogeny ^20^, external factors that affect the makeup of the human microbiome could have a measurable effect on the number of prophages. Human health is perhaps the most critical factor that influences the microbiome. Illnesses and generally poorer health are often associated with less diverse microbiomes, particularly in the respiratory and gastrointestinal systems ^1,2,38^.

Many of the samples were clinical samples which likely influenced the results, for instance, bacteria that dominate in dysbiotic microbiomes and specific disease-causing bacteria. To examine if these clinical samples exhibit different lysogenic profiles, we split the samples into groups based on whether the sample metadata listed the human as healthy, having various ailments (including diseases caused by specific bacteria and other ailments involving various bacteria/viruses), or asymptomatic. We independently assessed the samples by body site when investigating differences in the number of prophages per genome and determined significant differences using a Kruskal-Wallis test (Fig 3).

**Figure 3:**
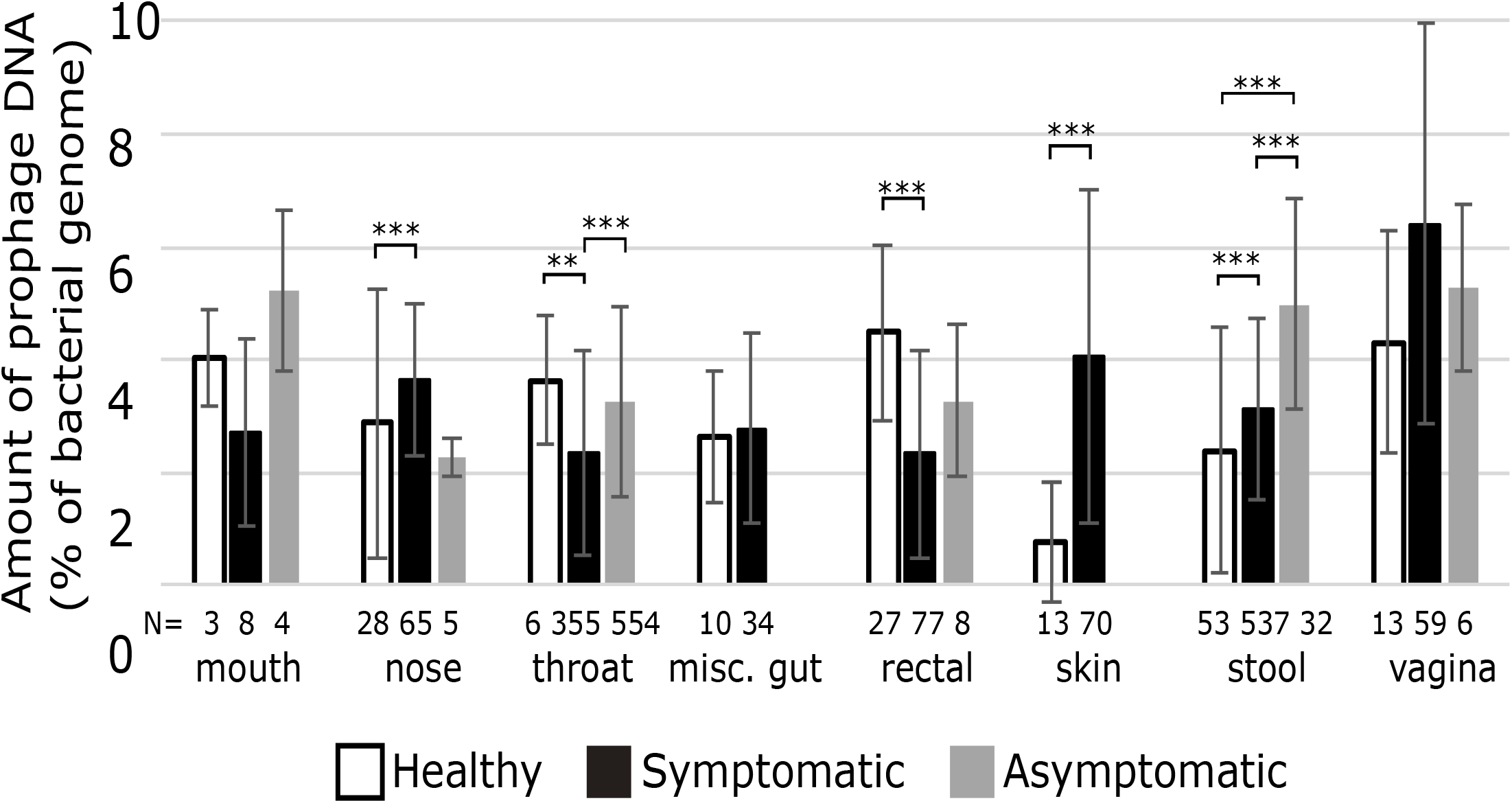
Each area of the human body with at least three genomes and described as sampled from healthy people. The number beneath each column indicates the number of samples in each group. Error bars represent one standard deviation, while asterisks represent significant differences (**= p=<0.005, ***= p=<0.001).

Only eight body sites had samples from healthy individuals (Fig. 3). The nose, skin, and stool samples all had more prophages in samples from symptomatic patients than healthy individuals, suggesting that prophages may contribute to disease at these sites. In contrast, the throat and rectal samples had fewer prophages in symptomatic individuals than in healthy people.

Both throat and stool samples had significantly more prophages in asymptomatic individuals than in healthy ones. Typically, patients are classified as asymptomatic when they suffer from a disease but are not currently experiencing symptoms. The difference in prophage abundance could suggest that prophages are either decreasing the virulence of their hosts in these areas or providing greater survivability so that once the illness clears, predominantly lysogens remain.

Related samples, such as from the lower gastrointestinal tract (stool and rectal) or the respiratory system (nose, mouth, and throat), did not always show similar patterns of prophage abundance. The bacterial species, types of illnesses, or the different types of tests used at different sites, could eliminate patterns between body sites.

### Conclusions

There is a lot of variation in the amount of prophage DNA in the bacteria of the human microbiome. Categorising the samples by body site revealed patterns in prophage abundance. Areas connected or with similar environments often had a similar prophage distribution. The respiratory tract, which has a lower microbial load, appears to follow the Piggyback-the-Persistent scenario, while the microbially rich gastrointestinal tract follows the Piggyback-the-Winner scenario. The microbiome impacts human health, and vice-versa, and a few body sites showed significant differences in prophage abundance in health and disease.

## Supporting information

Key resources table

## Author contributions

LKI generated all the data and figures and analysed all the data. MJR analysed the data and performed bioinformatics. RAE conceived the study. All three authors wrote the manuscript.

## Acknowledgements

Awards from the NIH NIDDK RC2DK116713 and the Australian Research Council DP220102915 to RAE supported this research. Flinders University Impact Seed Funding for Early Career Researchers supported MJR.

## Star Methods

### Resource availability

#### Lead contact

Laura K Inglis will answer and fulfil requests for further information and requests for resources and reagents.

## Materials availability

This study did not generate new unique reagents.

## Data and code availability

This paper analyses existing, publicly available data. The key resources table lists the accessions for these datasets.

This paper does not report any original code.

Any additional information required to analyse the data reported in this paper is available from the lead contact upon request

## Method details

All 949,935 publicly accessible genomes listed in the dataset “NCBI Genome Assemblies Summary Archive 20220601” (key resources) were downloaded from GenBank on June 1, 2022. PhiSpy ^39,40^ was used to analyse the genomes and detect prophage genomes in the bacterial DNA ^41^ as it is currently the best-performing prophage prediction tool ^42^. All the predicted prophages are available from FigShare (Key Resources “Prophage predictions”).

The data was filtered to remove any metagenome-assembled genomes, low-quality genomes with more than 50 contigs, duplicate genome sequences, and genomes not isolated from humans using the NCBI Genome Assemblies Summary (Key Resources “NCBI Genome Assemblies Summary Archive 20220601”). We manually sorted the remaining samples into categories and subcategories based on the area of the body from where they were isolated and the human host’s health according to the PATRIC metadata (Key Resources “Archive of the PATRIC Metadata from 20220601”) ^43^.

After filtering, 20,573 unique genome accessions remained. Over half, 11,513 genomes, came from bacteria associated with different human body areas. We separated those into 32 categories, with three to 2,970 samples per category. Approximately half of the genomes from human-associated bacteria, 6,844, could be categorised by the human host’s health. We provide this data as Key Resources “Prophages in humans”.

### Quantification and statistical analysis

SPSS was used for statistical analysis. The Kruskal-Wallis test was used to compare the categories of genomes associated with different human body areas.

We found bacterial concentrations for some of the categories by searching the literature. If we found multiple different estimates, we used an average of the concentrations to compare the prophage abundances with the bacterial concentrations.

For the genomes we could categorise by human health, we analysed multiple groups using SPSS. We compared the healthy, symptomatic, and asymptomatic groups within each category with either a Kruskal-Wallis test for categories with genomes in all three groups or Mann-Whitney-U tests for the skin and gut samples that only had two variables. We combined the categories and compared healthy, symptomatic, and asymptomatic groups with a Kruskal-Wallis test. Once we identified significant differences between healthy and symptomatic groups, we reanalysed the categories using only the healthy samples to determine whether the relative prophage abundances changed.

