## Supplementary material for "Prophage rates in the human microbiome vary by body site and host health": Key resources table

| REAGENT or RESOURCE | SOURCE | IDENTIFIER |
|---|---|---|
| Deposited data | | |
| NCBI Genome Assemblies Summary Archive 20220601 | (1) | https://doi.org/10.25451/flinders.22299664.v2 |
| Prophage predictions | (1) | https://doi.org/10.25451/flinders.c.6629843 |
| Archive of the PATRIC Metadata from 20220601 | (1, 2) | https://doi.org/10.25451/flinders.22299655.v2 |
| Prophages in humans | This paper | https://doi.org/10.25451/flinders.22758359.v1 |

### *References*


1. McKerral JC, Papudeshi B, Inglis LK, Roach MJ, Decewicz P, McNair K, Luque A, Dinsdale EA, Edwards RA. 2023. The Promise and Pitfalls of Prophages. bioRxiv.
2. Wattam AR, Abraham D, Dalay O, Disz TL, Driscoll T, Gabbard JL, Gillespie JJ, Gough R, Hix D, Kenyon R, Machi D, Mao C, Nordberg EK, Olson R, Overbeek R, Pusch GD, Shukla M, Schulman J, Stevens RL, Sullivan DE, Vonstein V, Warren A, Will R, Wilson MJC, Yoo HS, Zhang C, Zhang Y, Sobral BW. 2014. PATRIC, the bacterial bioinformatics database and analysis resource. Nucleic Acids Res 42:D581-91.
